# Intrinsic epigenetic state of primary osteosarcoma drives metastasis

**DOI:** 10.1101/2023.11.09.566446

**Authors:** Irtisha Singh, Nino Rainusso, Lyazat Kurenbekova, Bikesh K. Nirala, Juan Dou, Abhinaya Muruganandham, Jason T. Yustein

## Abstract

Osteosarcoma (OS) is the most common primary malignant bone tumor affecting the pediatric population with high potential to metastasize to distal sites, most commonly the lung. Insights into defining molecular features contributing to metastatic potential are lacking. We have mapped the active chromatin landscapes of OS tumors by integrating histone H3 lysine acetylated chromatin (H3K27ac) profiles (n=13), chromatin accessibility profiles (n=11) and gene expression (n=13) to understand the differences in their active chromatin profiles and its impact on molecular mechanisms driving the malignant phenotypes. Primary OS tumors from patients with metastasis (primary met) have a distinct active chromatin landscape compared to primary tumors from patients without metastatic disease (localized). The difference in chromatin activity shapes the transcriptional profile of OS. We identified novel candidate genes involved in OS pathogenesis and metastasis, including *PPP1R1B*, *PREX1* and *IGF2BP1*, which exhibit increased chromatin activity in primary met along with higher transcript levels. Overall, differential chromatin activity in primary met occurs in proximity of genes regulating actin cytoskeleton organization, cellular adhesion, and extracellular matrix suggestive of their role in facilitating OS metastasis. Furthermore, chromatin profiling of tumors from metastatic lung lesions noted increases in chromatin activity in genes involved in cell migration and key intracellular signaling cascades, including the Wnt pathway. Thus, this data demonstrates that metastatic potential is intrinsically present in primary metastatic tumors and the cellular chromatin profiles further adapt to allow for successful dissemination, migration, and colonization at the distal metastatic site.

## Introduction

Osteosarcoma (OS) is a primary malignant bone tumor commonly affecting children, adolescents, and young adults(1). In the last fifty years, the survival of patients with non-metastatic OS has significantly improved from approximately 20% to greater than 70%(2). However, approximately 20-25% of patients with OS present with metastatic disease and the outcomes for patients with metastatic or relapsed OS has remained unchanged over the past 30-40 years, with overall survival rates of less than 30%(2,3). Consequently, a better understanding of the molecular mechanisms that support metastasis is essential to improve the clinical outcome of patients with OS.

Enhancers are non-coding DNA regulatory elements typically bound by multiple transcription factors, mediators and RNA Poll complex that establishes a physical connection with the gene promoters to regulate cell type-specific gene regulatory profiles(4–7). Enhancer activity has been shown to be dysregulated in many cancers(8–13). A study that compared the epigenomic profile of metastatic lung OS tumors with primary bone OS tumors demonstrated positive selection of enhancer elements that endows the OS cells with metastatic competence(14). However, we still do not understand if some aspect of metastatic potential is intrinsically present in the primary bone tumor or is it secondary to acquired properties as the cells are presented with favorable conditions to migrate and colonize at a distant site. Here we report that primary bone tumors of OS patients that develop metastasis have both distinct chromatin activity and accessibility from the primary bone tumors of OS patients that do not present with metastatic disease. Thus, these intrinsic differences in the epigenetic state of primary tumor site shape the transcriptional profile of OS patients driving its metastatic fate.

## Materials and Methods

### PDX generation

Patient derived tumor xenograft (PDXs) were generated using tumor explants from biopsies and surgeries performed in patients with osteosarcoma. Tumors were established and propagated in the flank of 8-12 weeks old NOD-SCID-IL2γ-/- (NSG) mice as previously described(15). The generation harboring the patient-derived tumors was termed P0, with subsequent generations numbered consecutively P1, P2 and P3. Tumor histology confirmed the similarities between xenografts and original tumor specimens. The experiments described in the manuscript were performed using early passage (P1-P3) tumor generations.

### H3K27ac ChIP-seq sample and library preparation

The ChIP-seq experiment has been conducted by Diagenode ChIP-seq/ChIP-qPCR Profiling service (Diagenode Cat# G02010000). The chromatin was prepared by Diagenode ChIP-seq/ChIP-qPCR Profiling service (Diagenode Cat# G02010000) using the iDeal ChIP-seq kit for Histones (Diagenode Cat# C01010059). Chromatin was sheared using Bioruptor® Pico sonication device (Diagenode Cat# B01060001) combined with the Bioruptor® Water cooler for 8 cycles using a 30’’ [ON] 30’’ [OFF] settings. Shearing was performed in 1.5 ml Bioruptor® Pico Microtubes with Caps (Diagenode Cat# C30010016) with the following cell number: 1 million in 100μl. An aliquot of this chromatin was used to assess the size of the DNA fragments obtained by High Sensitivity NGS Fragment Analysis Kit (DNF-474) on a Fragment AnalyzerTM (Agilent). ChIP was performed using IP-Star® Compact Automated System (Diagenode Cat# B03000002) following the protocol of the aforementioned kit. Chromatin corresponding to 1 μg was immunoprecipitated using the following antibodies and amounts: 1 μg of H3K27ac (Diagenode C15410196; Lot A1723-00410). Chromatin corresponding to 1% was set apart as Input. qPCR analyses were made to check ChIP efficiency using KAPA SYBR® FAST (Sigma-Aldrich) on LightCycler® 96 System (Roche) and results were expressed as % recovery = 2^(Ct_input-Ct_sample).

The library preparation has been conducted by Diagenode ChIP-seq/ChIP-qPCR Profiling service (Diagenode Cat# G02010000). Libraries were prepared using IP-Star® Compact Automated System (Diagenode Cat# B03000002) from input and ChIP’d DNA using MicroPlex Library Preparation Kit v3 (Diagenode Cat# C05010001). Optimal library amplification was assessed by qPCR using KAPA SYBR® FAST (Sigma-Aldrich) on LightCycler® 96 System (Roche) and by using High Sensitivity NGS Fragment Analysis Kit (DNF-474) on a Fragment AnalyzerTM (Agilent). Libraries were then purified using Agencourt® AMPure® XP (Beckman Coulter) and quantified using QubitTM dsDNA HS Assay Kit (Thermo Fisher Scientific, Q32854). Finally, their fragment size was analyzed by High Sensitivity NGS Fragment Analysis Kit (DNF-474) on a Fragment AnalyzerTM (Agilent). Libraries were pooled and sequenced with Illumina technology with paired-end reads of 50bp length.

### ATAC-seq sample and library preparation

The ATAC-seq experiment was conducted by Diagenode ATAC-seq Profiling service (Diagenode Cat# G02060000). Tissues were weighted. 8.9-26.7 mg of tissue were chopped into small pieces (between 1-3 mm3) using a scalpel blade. Tissues were homogenized with a Dounce homogenizer and precleared by filtration through a 40μm Nylon cell strainer (Corning). Nuclei were then isolated on an iodixanol gradient and counted after addition of trypan blue using a TC20 automated cell counter (Biorad). 5.000-50.000 counted nuclei were transferred to a new tube and were centrifuged at 500g before proceeding with the transposition reaction. Isolated nuclei were lysed and transposed for 30 minutes at 37°c using the prokaryotic Tn5 transposase. Transposed DNA was then purified on Diapure columns (Diagenode, C03040001). The nuclei preparations were prepared using our standard protocol. The nuclei isolation was optimized on 11 mg of tissue. Two isolation conditions were tested: 0.1% Igepal CA-630 or 0.4% Igepal CA-630. The nuclei preparation was evaluated by visualization under the microscope (AE2000; Matic). We successfully isolated nuclei from Mouse PDX (Human) snap frozen tissues using 0.1% of Igepal CA-630 buffer. This condition was used to perform the ATAC assays on the samples of interest. Using the lysis conditions defined during the optimization, the nuclei were isolated from the different flash frozen tissue with a concentration of 0.1% of Igepal CA-630 in the lysis buffer. The purity of the nuclei isolation was evaluated by visualization under the microscope (AE2000; Matic) after the addition of trypan blue. Isolated nuclei were lysed and transposed for 30 minutes at 37°C using kit, Illumina, FC-121–1030). Transposed DNA was then purified on Diapure columns (Diagenode, Transposed purified DNA was then pre-amplified for 5 PCR Cycles using NEBNext High-Fidelity PCR MasterMix (NEB, M0541) and Illumina indexing primers. To define the final number of amplification cycles required, the pre-amplified libraries were analyzed by qPCR. After the amplification, the libraries were size selected then purified using AMPure beads and eluted in Tris pH7.5-8.0. The purified libraries were quantified using the Qubit ds DNA HS kit. The purified libraries were also analyzed on the Fragment Analyzer to assess their size. Using the quantification values from the Qubit and the size measurement generated by the Fragment Analyzer, the molar concentration of each library was calculated.

### RNA-sequencing Library Preparation and Sequencing

RNA was isolated using RNeasy (QIAGEN) and samples underwent quality control assessment using the RNA tape on Tapestation 4200 (Agilent) and were quantified with Qubit Fluorometer (Thermo Fisher). The RNA libraries were prepared and sequenced at the University of Houston Seq-N-Edit Core per standard protocols. RNA libraries were prepared with QIAseq Stranded Total RNA library Kit (Qiagen) using 500 ng input RNA. mRNA was enriched with Oligo-dT probes attached to Pure mRNA beads (Qiagen). RNA was fragmented, reverse transcribed into cDNA, and ligated with Illumina sequencing adaptors. The size selection for libraries was analyzed using the DNA 1000 tape Tapestation 4200 (Agilent). The prepared libraries were pooled and sequenced using NextSeq 500 (Illumina); generating ∼17million 2×76 bp paired-end reads per samples.

### H3K27ac ChIP-seq processing

The H3K27ac ChIP-seq was processed following the guidelines of ENCODE (phase-3) by using the ENCODE Transcription Factor and Histone ChIP-seq processing pipeline (https://github.com/ENCODE-DCC/chip-seq-pipeline2). This ChIP-seq processing pipeline (1) mapped the sequenced reads to the human reference genome hg38/GRCh38 using the Bowtie2, (2) filtered low-quality, duplicate, multimapping, unmapped reads along with reads mapping to the mitochondrial genome and the blacklisted regions, and (3) performed peak calling using MACS2 with a *P* value threshold of 10^−5^ (^16^). The quality control measures such as mapping statistics, library complexity (PCR bottlenecking coefficients, PBC1, and PBC2), cross-correlation scores (normalized strand cross-correlation coefficient and relative strand cross-correlation coefficient) and fraction of reads in the peaks (FRiP-score) as defined by ENCODE data standards were also determined using the pipeline(^16,17^). Additionally, we also determined the fraction of reads mapping within 2-kb of an annotated promoter as a quality control measure. We also aligned the reads to the mouse genome (mm10) and used XenofilterR to identify the ambiguous reads that align to both mouse and human genomes (18). This deconvolution process revealed that only 0.01-.04% reads align to both the genomes. The H3K27ac heatmaps and metaplots were generated using R Bioconductor package genomation(19).

### ATAC-seq processing

The chromatin accessibility sequencing data was processed following the guideline of ENCODE (phase-3) by using ENCODE ATAC-seq pipeline (https://github.com/ENCODE-DCC/atac-seq-pipeline). The ATAC-seq pipeline mapped the sequenced reads to the human reference genome hg38/GRCh38 using the Bowtie2, (2) filtered low-quality, duplicate, multimapping, unmapped reads along with reads mapping to the mitochondrial genome and the blacklisted regions, and (3) performed peak calling using MACS2 with a P value threshold of 10^−5^. We also aligned the reads to the mouse genome (mm10) and used XenofilterR to identify the ambiguous reads that align to both mouse and human genomes (18). This deconvolution process revealed that only 1.09-1.63% reads align to both the genomes. The chromatin accessibility heatmaps and metaplots were generated using R Bioconductor package genomation(19).

### RNA-seq processing

The sequenced reads of RNA-seq experiments were aligned to human reference genome hg38/GRCh38 using STAR aligner(20). The expression of the genes was determined as fragments per kilobase million (FPKM) by counting the number of uniquely mapping fragments and normalizing it by the length coding sequence and the library size. Gene annotation was based on UCSC RefSeq. Genes expressed with >1 FPKM in ≥50% of the samples of each group were considered to be expressed genes.

### DepMap data analysis

The CRISPR knockout screens data (22Q1) was obtained from the DepMap portal (https://depmap.org/)(21). The normalized dependency scores for all the genes and all the cell lines were utilized for the analysis. All the cancer cell lines associated with bone lineage were categorized as Bone while the remaining were referred as others.

### GO enrichment analysis

The R Bioconductor package clusterProfiler was used for gene ontology enrichment analysis(22). The genes that were < 50kb in distance of significantly affected H3K27ac peaks and were expressed with 1 FPKM in at least 50% of the samples of any of the three categories were utilized for the analysis. All the genes detected to be expressed from RNA-seq data were taken as the background. For the hierarchical clustering, first all the GO terms with significant enrichment were determined (*P* < 0.01, minimum gene set size > 10, maximum gene set size < 500), and then based on their semantic similarity were clustered into groups. For comparing the enriched biological themes across the groups, we first identified the significant terms (*P* < 0.001, minimum gene set size > 20, maximum gene set size < 300) and then compared them.

## Data availability

The data generated in this study are publicly available in Gene Expression Omnibus (GEO) at GSE234999.

## Results

### Primary tumors with impending metastasis exhibit distinct chromatin and epigenetic profiles

To investigate the differences in the active chromatin of OS, we utilized patient-derived xenografts (PDX) derived from tumor biopsies of patients with OS. These PDX models were generated using OS tumors obtained from either the primary bone site at diagnosis, refractory or recurrent tumors obtained from the primary site after surgery, or metastatic lung tumors (see Materials and Methods)(15). The PDX’s used in this study were grouped into the following categories based on disease presentation; **localized** - tumor obtained from the primary site in a patient without evidence of metastatic disease; **primary met** - tumor obtained from the primary site in patient with evidence of metastasis; **local control** - tumor obtained from primary site through surgical resection after neoadjuvant chemotherapy; and **distal met** - metastatic tumor obtained from a distal site (e.g., lung) (**Fig 1A**). Further clinical and pathological characteristics for these tumor specimens are presented in Supplementary Table 1.

**Figure 1:**
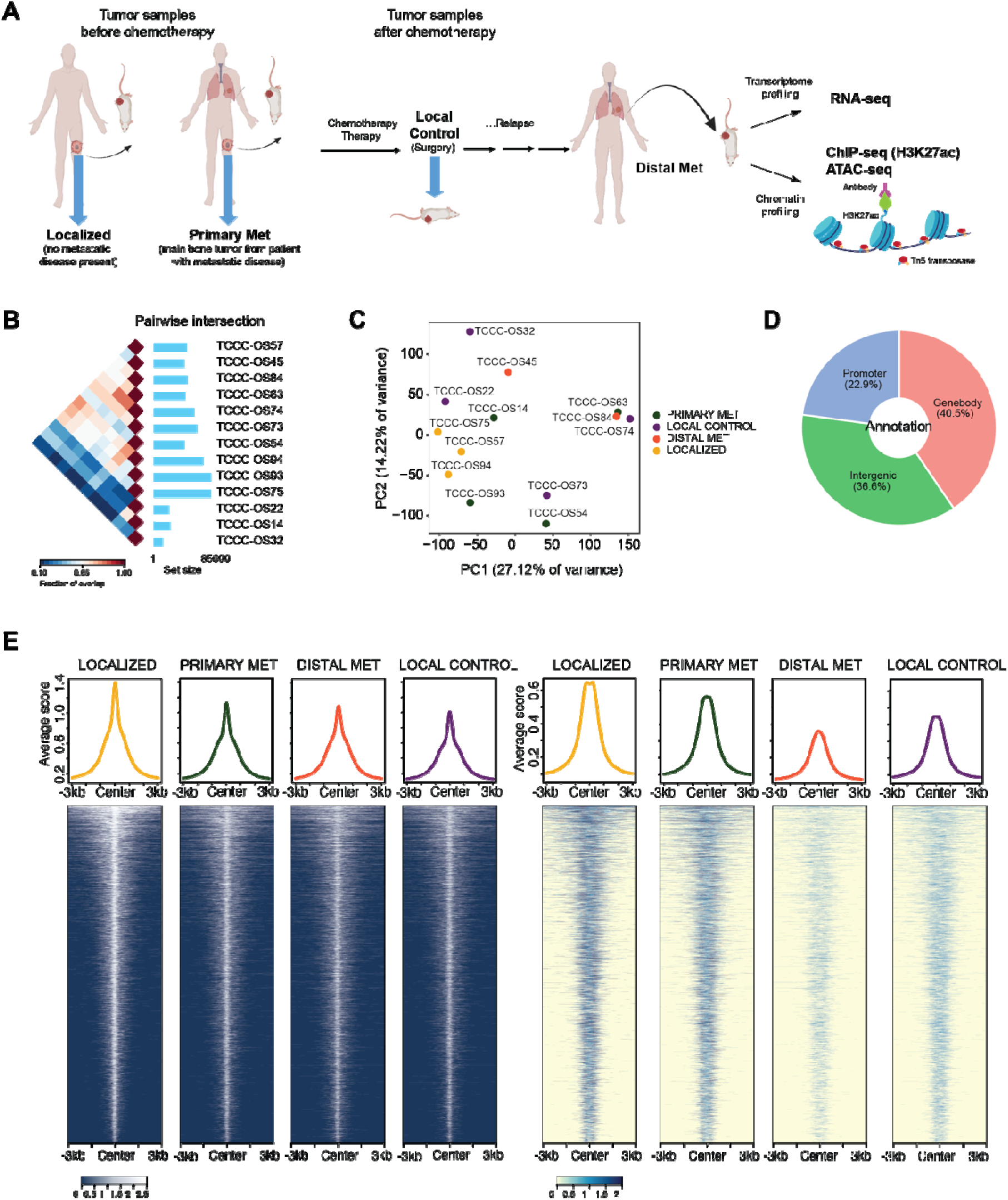
Active chromatin landscape of OS. **(A)** Schematic showing the generation of patient-derived xenografts (PDX) from primary OS tissues for each of the following categories; primary tumor obtained from the localized site that did not metastasize – **localized;** primary tumor obtained from the localized site that eventually underwent metastasis -**primary met**; tumor obtained from primary site after surgery and chemotherapy – **local control**; metastatic tumor obtained from a distal site – **distal met**. The PDX’s subsequently underwent transcriptomic by RNA-seq and epigenomic profiling by ATAC-seq and ChIP-seq for H3K27ac. **(B)** Shows the overlap between the H3K27ac peaks detected for each sample using intervene tools(52). **(C)** Principal component analysis of signal intensity (tags per million) of top 20% most variable regions by median absolute deviation (n = 31,690) of all H3K27ac enriched peaks. **(D)** Shows the fraction of H3K27ac enriched peaks (n = 158,455) overlapping with different genomic annotations. Peaks overlapping within ±LJ2.5KB of any annotated transcription start site were considered to be promoter associated peaks (22.90%); peaks overlapping with 5’UTR, coding exons, introns, and 3’UTRs were considered to be in gene bodies (40.50%) and all the remaining peaks were considered to be intergenic peaks (36.60%). **(E)** Left - Heatmaps of average H3K27ac activity ±LJ3LJKb relative to the center of the detected peak for localized (yellow), primary met (green), local control (red) and distal met tumors (purple), metaplots showing the average levels of H3K27ac on top each of the heatmap (top). These regions were identified to be enriched for H3K27ac signal in every sample (n = 6,086). Right - Heatmaps of average chromatin accessibility at the corresponding locus for every row in D, metaplots showing the average chromatin accessibility on top each of the heatmap (top)

We integrated active chromatin landscapes defined by histone H3 lysine acetylated chromatin (H3K27ac) and chromatin accessibility profiled by assay for transposase-accessible chromatin (ATAC) with gene expression across a cohort of localized (n = 3), primary met (n = 4), local control (n = 4) and distal met (n = 2) PDX’s (Supplementary Table 2, See Materials and Methods). Amongst our PDX models, TCCC-OS63 (primary met), TCCC-OS74 (local control tumor with poor response to chemotherapy) and TCCC-OS84 (lung metastasis) were generated from tumor tissue obtained from one individual patient at different phases of the disease progression. TCCC-OS54 (primary met) and TCCC-OS73 (local control with poor response to chemotherapy) were also generated using tissues from the same patient. Thirteen PDXs were profiled by at least one genomic readout, and within this dataset, 11 samples were matched for H3K27ac, chromatin accessibility, and gene expression (**Fig 1A**). Every H3K27ac ChIP-seq sample was accompanied with background ChIP-seq with no specific pull-down. H3K27ac ChIP-seq and ATAC-seq datasets were subjected to rigorous quality control based on Encyclopedia of DNA elements (ENCODE) best practices(23). We used ENCODE (phase-3) ChIP-seq and ATAC-seq pipeline to identify genomic regions with significantly enriched H3K27ac levels and chromatin accessibility, respectively (See Materials and Methods).

There was high pairwise overlap between enriched H3K27ac regions of all PDX models except TCCC-OS14, 22 and 32 (**Fig 1B**). The reduced overlap of H3K27ac enriched regions of TCCC-OS14, 22 and 32 with other samples was driven by lower quality of H3K27ac ChIP-seq experiments (low FRiP scores and low peak detection) rather than real differences in their epigenomic profile (Supplementary Table 3). To work with robust H3K27ac enriched genomic regions, we filtered for regions that were identified in at least two PDXs (n = 158,455). Comparison of signal intensity across top 20% most variable H3K27ac regions (n = 31,690) revealed OS tumors obtained from the same patient to exhibit higher similarity to each other despite having distinct clinical and pathological features (**Fig 1C, S1**). Overlap of H3K27ac enriched regions with genomic annotation revealed that majority of these regions are enhancer elements lying further away from the gene promoters (**Fig 1D**). Peaks overlapping within ±LJ2.5KB of any annotated transcription start site were considered to be promoter associated peaks, peaks overlapping with 5’ untranslated regions (5’UTR), coding exons, introns and 3’ untranslated regions (3’UTRs) were considered to be in gene bodies and all the remaining peaks were considered to be intergenic peaks. Almost a quarter of H3K27ac enriched regions overlapped with gene promoters, an observation that is in concordance with established standards of H3K27ac profiling. The chromatin activity and corresponding accessibility at regions that had significant H3K27ac enrichment in every PDX sample further validates the rigor of our epigenomic profiling (**Fig 1E**).

To delineate epigenetic features that potentially play a role in defining the metastatic potential of the primary tumor, we compared the epigenetic, chromatin, and transcriptional profile of primary met and localized tumors. We utilized generalized linear models to identify the significantly differential H3K27ac regions between primary mets and localized tumors(24). Differential analysis of H3K27ac regions revealed 4,099 (*P-adjust* < 0.2; *P* < 0.0071) genomic regions that exhibit distinct extent of chromatin activity in primary met and localized tumors (**Fig 2A**). When we overlapped the differential H3K27ac regions with genomic annotation, we found that majority of these regions were in gene bodies or intergenic regions (**Fig 2B**). This observation suggests that most of the differences in chromatin activity between primary mets and localized tumors occurs at enhancer elements. Briefly, 2,219 genomic regions showed enhanced chromatin activity in primary met, while 1,880 genomic regions showed reduced chromatin activity in comparison to localized tumors (**Fig 2C-D**). The corresponding chromatin accessibility to the differential H3K27ac regions corroborates these differences in chromatin activity between primary met and localized tumors. To evaluate the effect of the differences in chromatin activity on transcriptional output, we also compared the mRNA levels of the proximal genes. The genes proximal to regions with enhanced chromatin activity overlapping with different genomic annotations consistently had higher mean mRNA expression levels in primary met compared to localized tumors (**Fig 2E**; genes (n=207) with differential chromatin activity overlapping with promoters - *P* < 9.26 x 10^-33^ by Wilcoxon signed rank test; genes (n = 360) with differential chromatin activity overlapping with gene bodies - *P* < 6.28 x 10^-39^ by Wilcoxon signed rank test and genes (n = 188) with differential chromatin activity overlapping with intergenic regions *P* < 2.74 x 10^-17^ by Wilcoxon signed rank test). Similarly, genes with reduced chromatin activity in primary met compared to localized tumors had lower expression levels (**Fig 2E**; genes (n=121) with differential chromatin activity overlapping with promoters - *P* < 9.85 x 10^-21^ by Wilcoxon signed rank test; genes (n = 410) with differential chromatin activity overlapping with gene bodies - *P* < 2.24 x 10^-46^by Wilcoxon signed rank test and genes (n = 160) with differential chromatin activity overlapping with intergenic regions 9.74 x 10^-21^ by Wilcoxon signed rank test). As expected, the extent of this difference in mRNA levels was more for promoter associated elements in comparison to gene bodies and intergenic regions. When we compared only the transcript levels between primary met and localized tumors, fewer genes (n=192) were noted to be significantly differentially expressed (*P-adjust < 0.05*) **(Fig S2).** Thus, differences in the active chromatin profiles provides the ability to identify a more comprehensive repertoire of significant, and potentially biologically relevant genetic changes that can go undetected if only analyzing transcript levels. Overall, these data suggest that the enhancer landscape of primary metastatic tumors is distinct from primary localized tumors and significantly impacts the transcriptional output.

**Figure 2:**
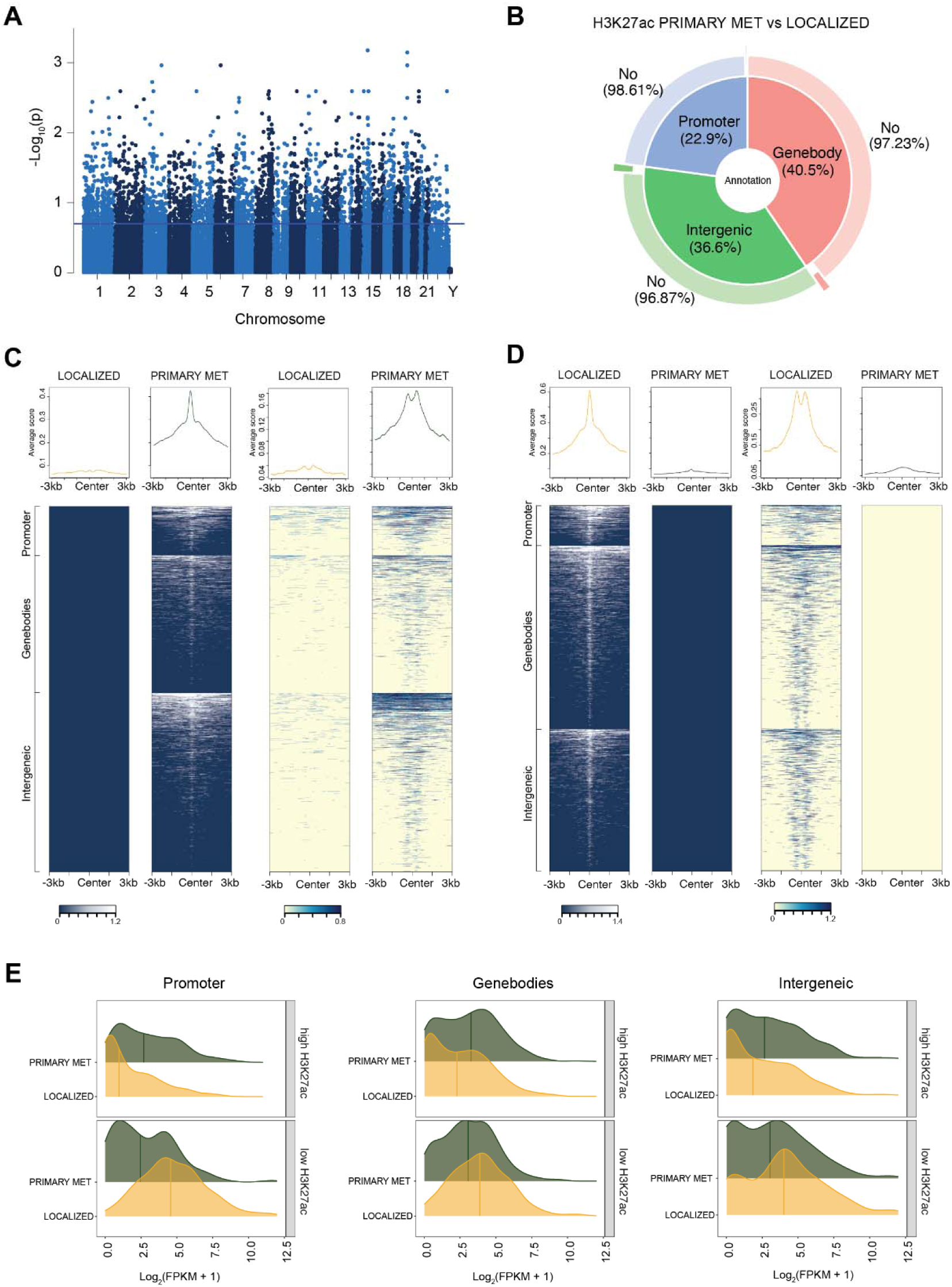
Differential chromatin activity between localized and primary met. **(A)** Manhattan plots showing the regions with differential H3K27ac activity (*P-adjust* < 0.2 corresponding to *P* < 0.0071) between primary met and localized tumors across chromosomes. **(B)** As in Figure 1C, but also showing the fraction of peaks overlapping with each of the regions that show differential H3K27ac activity. 1.39% of all promoter peaks, 2.77% of peaks overlapping with gene bodies, and 3.13% of intergenic peaks show significant differential activity between primary met and localized tumors. **(C)** Left, as in figure 1D for peaks with high H3K27ac activity in primary met compared to localized tumors. Top is for peaks lying in the promoter region (n = 300), middle is for peak overlapping with gene bodies (n = 833) and bottom is for peaks overlapping with intergenic regions (n = 1,086), metaplots showing the average levels of H3K27ac are shown on top each of the heatmap (top); Right, as in figure 1E, shows the chromatin accessibility at the locus for every corresponding row in figure 2C **(D)** As in Figure 2C for peaks with low H3K27ac activity in primary met compared to localized tumors. Top is for peaks that lie in the promoter region (n = 205), middle is for peak overlapping with gene bodies (n = 944) and bottom is for peaks overlapping with intergenic regions (n = 731), metaplots shows the average levels of H3K27ac are shown on top each of the heatmap (top); Right, as in figure 1E, shows the chromatin accessibility at the locus for every corresponding row in figure 2C. **(E)** Density plot of mean RNA expression levels of genes proximal (< 50Kb) to differential H3K27ac peaks in primary met and localized tumors. Top to bottom shows the following order, genes with high H3K27ac levels in promoters of primary met (n = 207, *P* < 9.26 x 10^-33^ by Wilcoxon signed rank test), genes with low H3K27ac in promoters of primary met (n = 121, *P* < 9.85 x 10^-21^ by Wilcoxon signed rank test), genes with high H3K27ac levels in gene bodies of primary met (n = 360, *P* < 6.28 x 10^-39^by Wilcoxon signed rank test), genes with low H3K27ac in promoters of primary met (n = 410, *P* < 2.24 x 10^-46^ by Wilcoxon signed rank test), genes with high H3K27ac levels in intergenic regions of primary met (n = 188, *P* < 2.74 x 10^-17^ by Wilcoxon signed rank test) and genes with low H3K27ac in intergenic regions of primary met (n = 160, *P* < 9.74 x 10^-21^ by Wilcoxon signed rank test).

### Enhancer activity shapes the gene expression profiles of primary tumors to facilitate metastasis

Next, we wanted to determine the genes whose expression was potentially affected by the differential enhancer activity and further investigate their role in OS pathogenesis and metastasis. We observed increased enhancer activity around many gene loci that could enable the primary met increased metastatic potential, thus permitting their migration and metastasis. Some of these candidates include *PPP1R1B*, *PREX1* and *IGF2BP1* (**Fig 3**). Phosphoprotein phosphatase-1 regulatory subunit 1B (PPP1R1B) encodes a bifunctional signal transduction protein, commonly referred as DARPP-32 (Dopamine and cyclic adenosine monophosphate-regulated phosphoprotein, Mr 32000). Its phosphorylation is regulated by dopaminergic and glutamatergic receptor stimulation. DARPP-32 has been found to be amplified or overexpressed in gastric, colon, prostate, and breast cancers(25–30). Reports have implicated DARPP-32 in cancer cell proliferation, survival, invasion and angiogenesis(31). More recently, a study showed the role of DARPP-32 in promoting cell survival and migration through non-canonical NF-κB2-p52 pathway in lung cancer(32). To identify if PPP1R1B confers the cancer cells with specific vulnerabilities, we utilized the genome-wide genetic screen for identifying cancer dependencies provided through the DepMap project(21). Compared to all the other genes, PPP1R1B showed a significant selective dependency in bone sarcomas as well as other cancers (**Fig 3C**; *P* < 8.47 x 10^-8^ by one-sided Kolmogorov-Smirnov test for PPP1R1B gene effect in bone sarcomas vs all other genes; *P* < 2.2 x 10^-16^ by one-sided Kolmogorov-Smirnov test for PPP1R1B gene effect in bone sarcomas vs all other genes). To our knowledge the role of DARPP-32 in OS metastasis remains unexplored. Our observation of increased chromatin activity, accessibility accompanied by increased mRNA levels of PPP1R1B, and selective dependency in primary mets warrants it further investigation. IGF2BP1 is shown to be the post-transcriptional regulator of mRNA targets that control cell proliferation and growth, invasion, and chemo-resistance(33,34). It is also associated with a poor overall survival and metastasis in various types of human cancers. However, its role in driving OS metastasis has not been determined. Strikingly, bone sarcomas exhibit significant selective survival dependency on IGF2BP1 when compared to all the other cancers **(Fig 3C**; *P* < 1.63 x 10^-8^ by one-sided Kolmogorov-Smirnov test for IGF2B1 gene effect in bone sarcomas versus other cancers).

**Figure 3:**
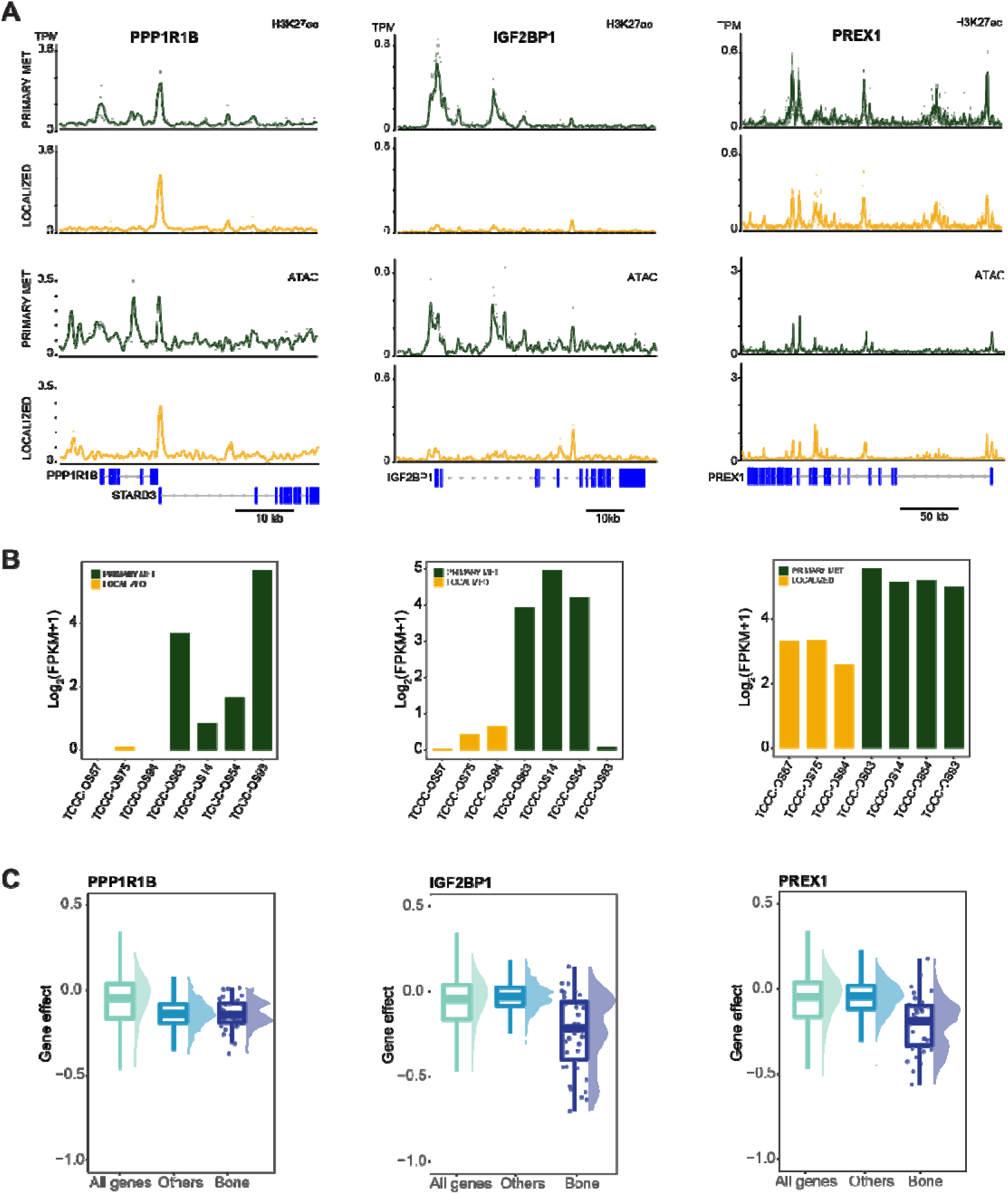
Enhancer activity shapes distinct gene expression profiles. **(A)** Top, H3K27ac activity at PPP1R1B, PREX1 and IGF2BP1 genomic loci in localized and primary met tumors. Bottom, chromatin accessibility as determined by ATAC-seq at PPP1R1B, IGF2BP1 and PREX1 genomic loci in localized and primary met tumors. The y-axis shows tags per million (tpm) for H3K27ac and chromatin accessibility levels for every sample. The thickened line is the average profile across all samples. **(B)** mRNA expression levels of *PPP1R1B, IGF2BP1 and PREX1* as determined by RNA-seq. **(C)** Genetic dependency of bone related sarcomas and other cancers on PPP1R1B (*P* < 8.47 x 10^-8^ by one sided Kolmogorov-Smirnov for PPP1R1B gene effect in bone sarcomas vs all other genes; *P* < 2.2 x 10^-16^ by one sided Kolmogorov-Smirnov for PPP1R1B gene effect in bone sarcomas vs all other genes), IGF2BP1 (*P* < 1.63 x 10^-8^ by one side Kolmogorov-Smirnov for IGF2B1 gene effect in bone sarcomas versus other cancers) and PREX1 (*P* < 1.7 x 10^-8^ by one side Kolmogorov-Smirnov for PREX1 gene effect in bone sarcomas versus other cancers) as measured by Cancer Dependency Map (DepMap) project. DepMap provides selective dependency score for every gene as determined by genome-scale CRISPR–Cas9 loss-of-function screens of 18,333 genes in 769 cell lines.

Like PPP1R1B and IGF2BP1, we found high chromatin activity around PREX1 locus with increased mRNA levels **(Fig 3A, 3B).** P-REX1 is a guanine nucleotide exchange factor that positively regulates Rac1-mediated oncogenic signaling(35–39). A recent report showed that in murine models, P-Rex1 cooperates with the neu oncogene to increase mammary tumor incidence and metastasis, but not primary tumor growth(40). Several studies also show PREX1 to be amplified in gynecological cancers **(Fig S3)**. Interestingly, bone sarcomas had selective genetic dependency on PREX1 compared to other cancers **(Fig 3C**; *P* < 1.7 x 10^-8^ by one-sided Kolmogorov-Smirnov test for PREX1 gene effect in bone sarcomas versus other cancers). Our data demonstrates how chromatin activity can regulate the gene expression of oncogenes in OS tumors.

We next investigated the biological process and molecular functions of all the genes proximal to regions with differential H3K27ac activity. Gene Ontology (GO) enrichment analysis of the proximal genes showed significant enrichment for biological processes involved in cell adhesion and migration (**Fig 4A**). Further, the GO analysis for molecular functions highlighted significant enrichment for genes encoding for proteins that are components of extracellular matrix, actin cytoskeleton, cell anchoring junction and actomyosin (**Fig 4B**). Thus, our results indicate that the primary met had differential chromatin activity around genes that regulate actin cytoskeleton organization, cellular adhesion, and migration. This difference in epigenetic state in primary met is suggestive of its role in facilitating OS metastasis.

**Figure 4:**
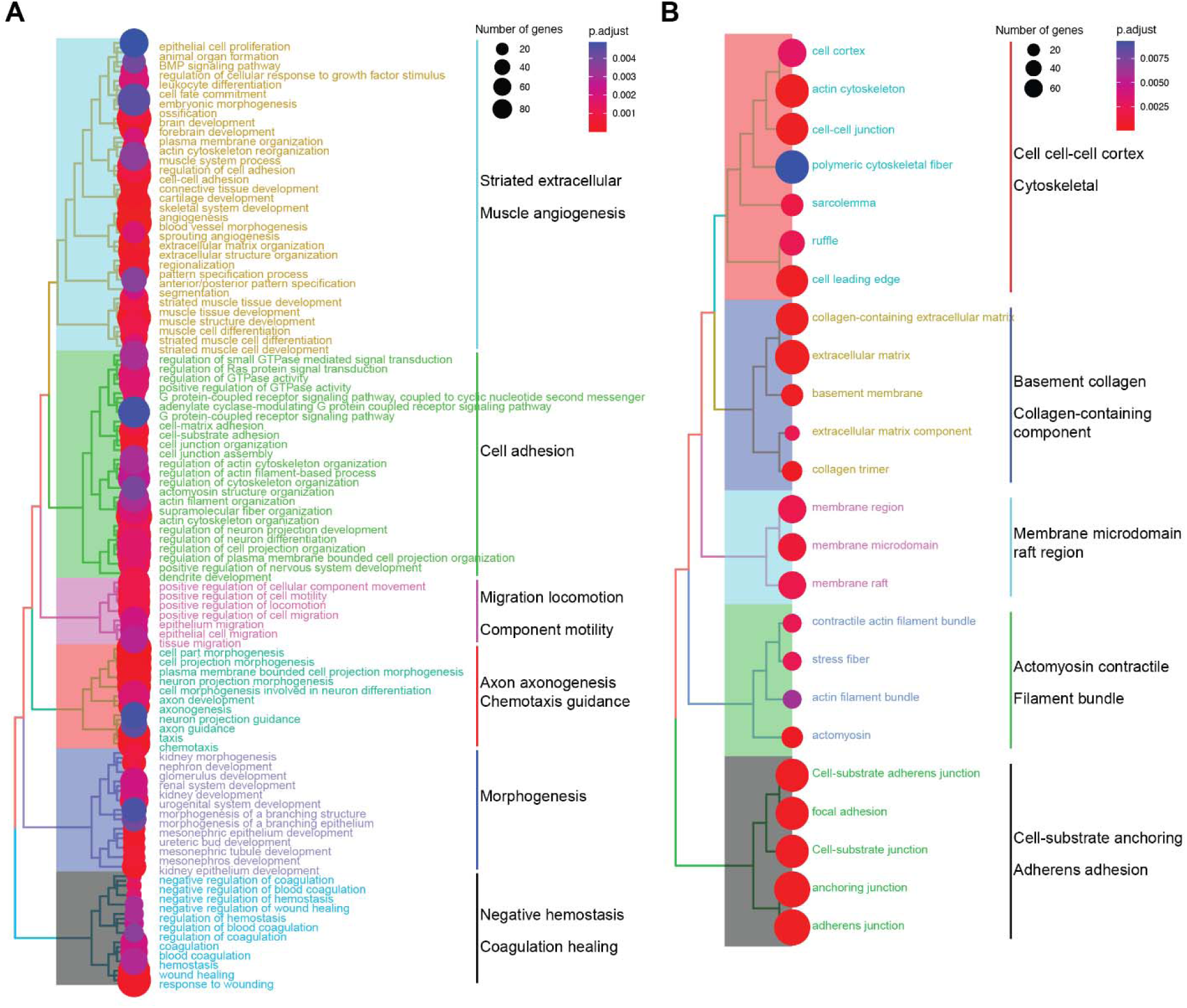
Molecular processes affected by chromatin profiles. **(A)** Biological process enrichment of genes proximal to differential H3K27ac regions (< 50Kb) between primary met and localized tumors. **(B)** As in (A) but for molecular functions (see Materials and Methods).

### Genome-wide differences in chromatin accessibility between primary met and localized tumors

Given the significant differences in the enhancer landscape of primary met in comparison to localized tumors, which reflect the chromatin accessibility of those genomic regions, we wanted to identify the transcription factors that were potentially occupying those accessible regions. All the accessible regions that were present in at least two samples (n = 121,024) were subjected to differential analysis using generalized linear models. We identified genomic regions that had higher (n = 3,662, *P-adjust* < 0.2 that corresponded to *P* < 0.02) and lower (n = 7,936, *P-adjust* < 0.2 that corresponded to *P* < 0.02) accessibility in primary met compared to localized tumors (**Fig 5A**). To identify the transcription factors that potentially occupy the differentially accessible regions, we performed HOMER motif analysis of highly accessible regions with lowly accessible regions as the background(41). This approach revealed that the highly accessible regions in primary met were enriched with binding motifs for the E2FA, EBF1/2/3, DLX1, HIC1/2, NFIX transcription factors. We repeated the motif analysis of lowly accessible regions in primary met with highly accessible regions as the background. We found that these regions with low accessibility in primary met that were open and accessible in localized tumors had significant enrichment for binding sites of AP1 (JUN/FOS) and TEAD family of transcription factors (**Fig 5B)**. Interestingly, EBF2 has previously been implicated in regulating metastatic and anti-apoptotic properties in osteosarcoma, while DLX1 and NFIX have been recently reported to be critical transcription factors in prostate cancer and lung cancer metastasis, respectively(42–45). There are no studies reporting the role of DLX1 and NFIX in OS metastasis, thus our chromatin profiling of these different states has identified potential new candidate transcriptional networks critical for metastatic OS.

**Figure 5:**
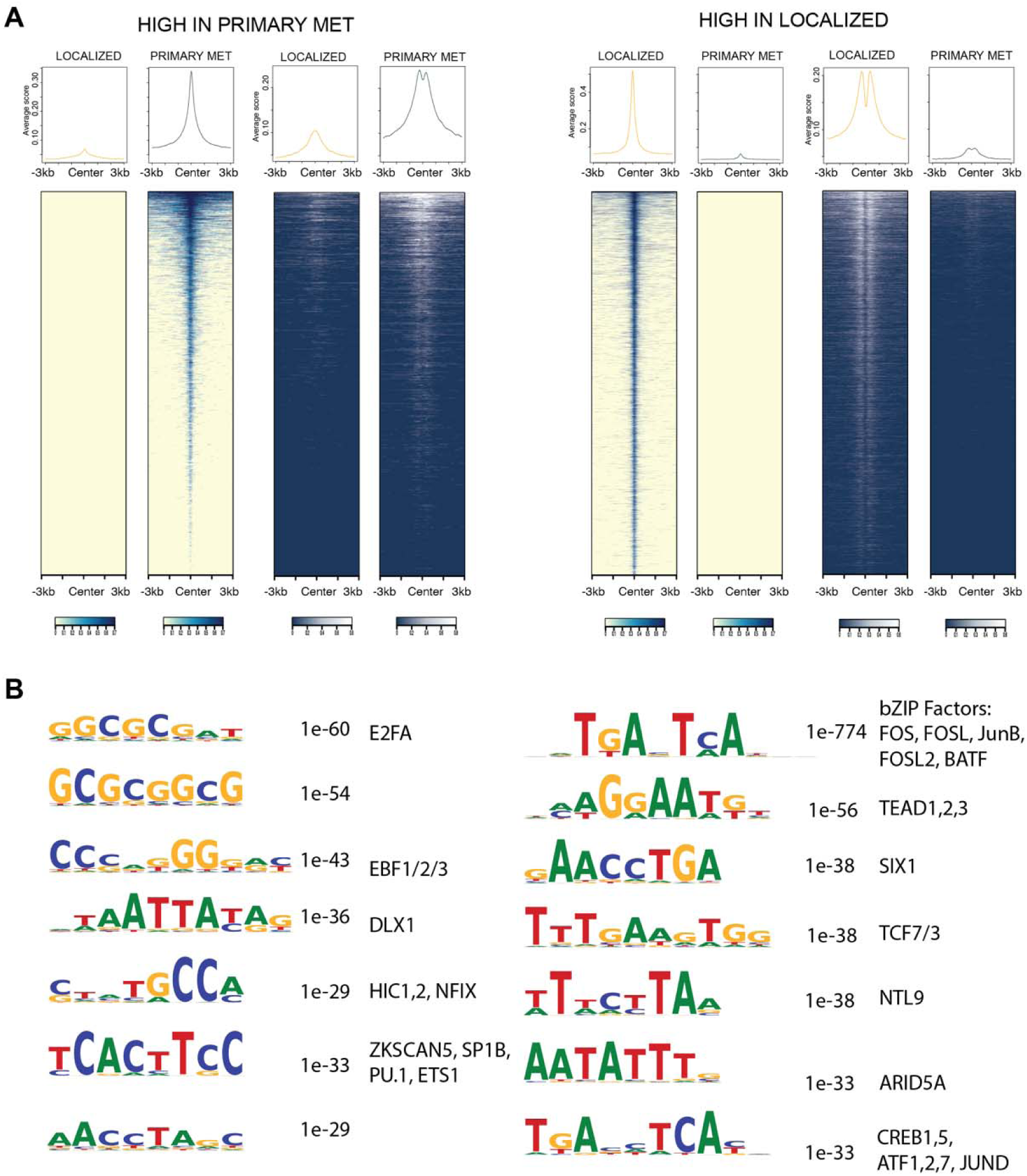
Differential chromatin accessibility between primary met and localized tumors. **(A)** Left, as in figure 1D for peaks with high chromatin accessibility in primary met compared to localized tumors (n = 3,662, *P-adjust* < 0.2 that corresponded to *P* < 0.02) with corresponding H3K27ac activity at the same loci, metaplots showing the average levels of chromatin accessibility and H3K27ac are shown on top each of the heatmap (top). Right, as in figure 1D, but for peaks with low chromatin accessibility in primary met compared to localized tumors (n = 7,936, *P-adjust* < 0.2 that corresponded to *P* < 0.02). (**B**) Left, sequence motifs enriched in the highly accessible chromatin region of primary met compared to localized tumors (lowly accessible regions were used as the background); Right, sequence motifs enriched in the lowly accessible chromatin region of primary met which exhibit high accessibility in localized tumors (highly accessible regions were as the background).

### Chromatin activity of distal met differs further from localized tumors compared to primary met

Given the observed differences in chromatin activity of primary met when compared to localized tumors, we investigated for differences in chromatin activity in the distal met and local control tumors. Differential analysis of H3K27ac regions (n = 158,455) between distal met and localized tumors as well as local control and localized tumors revealed 6,177 (*P-adjust* < 0.2 corresponding to *P* < 0.011) and 5,486 (*P-adjust* < 0.2 corresponding to *P* < 0.007) differential genomic regions, respectively. Out of 6,177 differential genomic regions, 3,308 regions had higher chromatin activity while 2,869 genomic regions displayed reduced chromatin activity in distal met when compared to localized tumors. Similarly, we identified 3,501 genomic regions with higher chromatin activity and 1,985 genomic regions with diminished chromatin activity in local control tumors when compared to localized tumors. The differences in the chromatin activity significantly impacts the transcriptional output of proximal genes (**Fig S4**).

Next, we wanted to determine if these regions are the same ones that we had determined to be differential in primary met when compared to localized tumors. Interestingly, only 505 genomic regions had significantly higher chromatin activity in all the three categories while 1,005, 1,752 and 1,762 genomic regions were detected to be significantly higher only in primary met, distal met and local control tumors, respectively (**Fig 6A**). We detected 263, 446 and 788 genomic regions with higher chromatin activity only in primary met and distal met, only in primary met and local controls, and only in distal met and local controls respectively. Doing a similar analysis for genomic regions with reduced chromatin activity, 278 genomic regions were significantly different in all three categories while 945, 1,789 and 992 regions were significantly differential only primary met, distal met and local controls, respectively (**Fig S5**). 372, 285 and 430 genomic region displayed lower chromatin activity only in primary met and distal met, only in primary met and local controls, and only in distal met and local controls, respectively.

**Figure 6:**
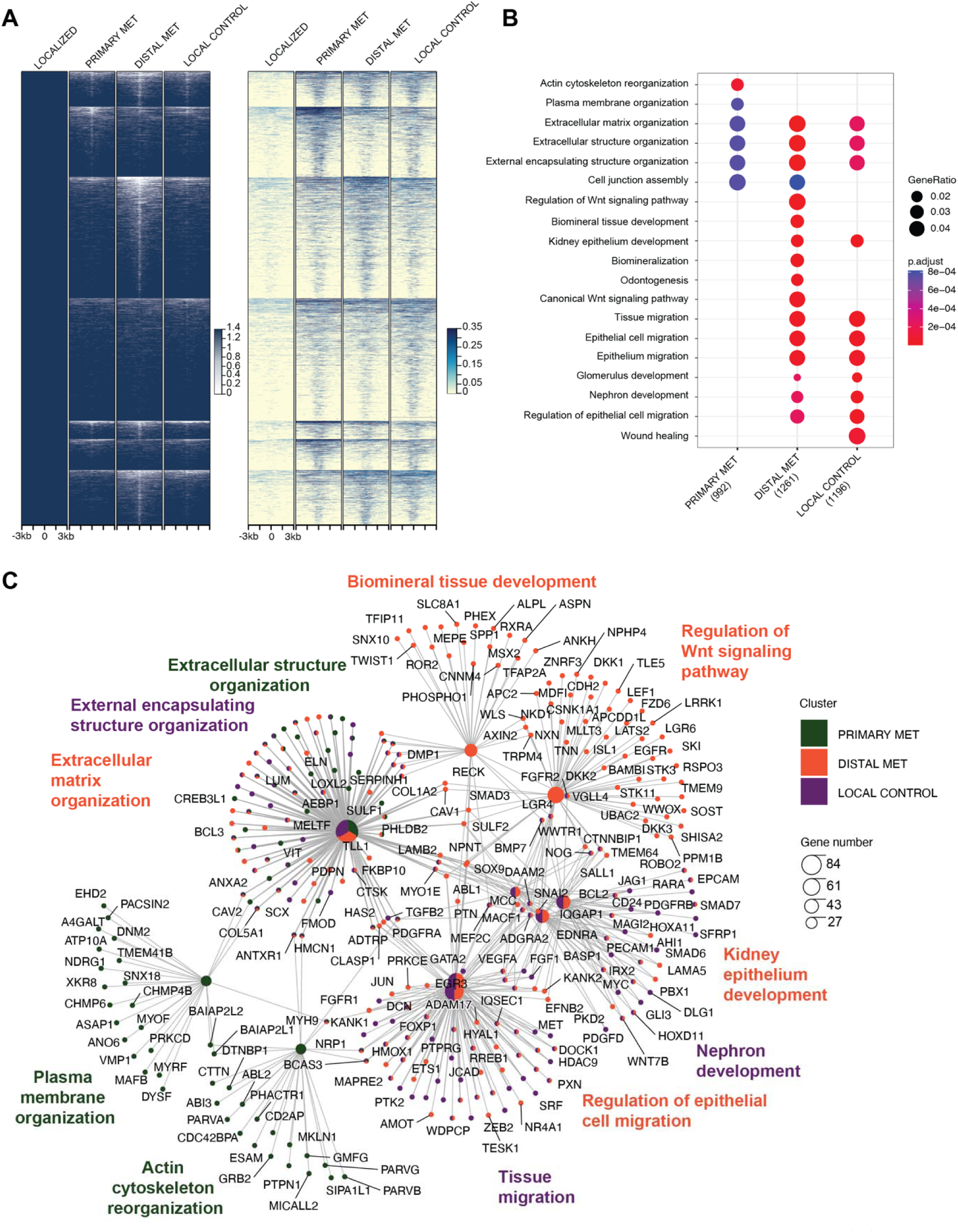
Distal met tumors exhibit further differences in chromatin activity. **(A)** Left, as in figure 1D but for peaks with high H3K27ac activity in primary met, distal met and local control (n = 505) when compared to localized tumors. Top is for peaks that had significantly high H3K27ac levels in each primary met, distal met and local control. It is followed by regions that had significantly high H3K27ac levels only in primary met (1,005), only in distal met (1,752) and only in local control (1,762). Below are genomic regions with high H3K27ac levels in primary met and distal met (263), primary met and local control (446) and distal met and local control (788). **(B)** Comparison of top biological processes enrichment of genes proximal to differential H3K27ac regions (< 50Kb) between each of the three categories (*P* < 0.01) **(C)** Highlighting proximal genes involved in the enriched biological processes shown in B.

From these findings we inferred that the distal met exhibit further changes in chromatin activity in comparison to what we detected in primary met relative to localized tumors. This differential chromatin activity in distal met might be responsible for shaping the gene expression dynamics allowing the tumor to migrate, establish and grow at a distal site. We acknowledge that these numbers are affected by significance cutoffs as well as sample sizes and thus should be interpreted cautiously. To examine the genes proximal to regions of differential chromatin activity and identify the biological processes they are involved in, we compared the top GO biological processes terms (*P* < 0.01) enriched in primary met, distal met and local control. We found Wnt signaling pathway to be affected in distal met. Wnt signaling is well known for its role in regulating key cellular functions including proliferation, differentiation, migration, genetic stability, apoptosis, and stem cell renewal. There have been reports implicating the role of aberrant Wnt signaling for OS growth, colony formation, invasion, and metastasis(46). Additionally, we also found biological processes related to cell migration to be enriched in the top terms in distal met and local control but not in primary met tumors. Interestingly, actin cytoskeleton reorganization was enriched in primary met, but not in distal met and local control tumors. **Figure 6B** highlights the genes involved in these biological processes and categories they were enriched in. In summary, these findings suggest the distal metastatic tumors undergo further changes in its epigenomic profile to eventually migrate and colonize.

## Discussion

The successful treatment of metastatic disease remains an extremely difficult challenge for solid tumor patients, including osteosarcoma. The identification of key underlying molecular mechanisms responsible for predisposing and enabling the tumor cell to enhance its metastatic potential remains elusive. Using PDX specimens, we integrated epigenetic analysis with transcriptomic profiling from patients with OS diagnosed with upfront metastatic and localized disease and identified novel differential genetic features that are candidate innate metastatic-promoting genes. We understand that while using PDX specimens obtained from subcutaneous engraftment grown in immunodeficient mice can cause some potential changes in tumor intrinsic profiles, our approach of placing 1-2mm slices of often minimal biopsied tumor quantity allows for retention of the cellular heterogeneity and architecture while also providing extremely valuable additional tumor specimen for additional downstream molecular and therapeutic studies upon PDX growth. Unfortunately, this approach does allow orthotopic placement of the tumors since direct injection into the tibia or tail vein (to mimic metastasis) of the tissue slices is not feasible. However, there have been several prior reports validating the use of subcutaneous PDXs for preserving genetic profiling and also their high therapeutic predictive potential (47) (48) (49) (50) (51).

Through integration of RNA-Seq, ATAC-seq, and ChIP-Seq for H3K27Ac we identified significant differential epigenomic landscapes for primary tumors obtained from patients with localized and metastatic OS, with noted differential H3K27Ac activity across the genome for those patients with upfront metastatic disease. We have stratified the patients into these categories based on their radiographic staging at time of diagnosis, which still has high prognostic significance. We understand that untreated localized patients would still have high chance of relapse without systemic chemotherapeutic intervention, but also recognize that upfront staging does have clinical and conceptually molecular significance for more advanced, metastatic disease. Through these studies we are attempting to identify intrinsic molecular signatures that can be contributing to the more advanced upfront disease.

Our identification of novel candidate metastatic-promoting genes that are present in primary metastatic bone tumors, such as PPP1R1B, PREX1 and IGF2BP1, provides the framework for subsequent functional genomic studies into the role of these genes in metastatic osteosarcoma. Specifically, PPP1R1B, referred to as DARPP-32, encodes a bifunctional signal transduction protein involved in multiple carcinomas such prostate, colon and breast and is involved in proliferation, survival, invasion, and angiogenesis. Our data validates the role of PREX1 as an oncogene in OS primary met tumors.

Overall, there has been a paucity of investigations into the epigenetic profiles involved in metastatic osteosarcoma. Morrow et al. profiled paired primary and lung metastatic tumors from the same patient (5 patients) as well as paired human cell lines (e.g., HOS/143B)(14). Their analysis of these tumors and cell lines using ChIP-seq for H3K4me1, H3K27Ac identified Met-variant enhancer loci (VEL) distribution clusters at distinct regions across the epigenome that were often in the vicinity of individual genes. Though this manuscript provided valuable insights into distal lung metastasis enhancer profiles, our study is focused on the underlying promoter and enhancer profiles between the primary tumors from localized and metastatic patients, which confer innate metastatic-promoting mechanisms that can be responsible for driving metastasis from the originating bone tumor. In addition, our study has integrated RNA-seq analysis to identify key gene ontology sets that provide a more global mechanistic understanding of tumor dissemination. Finally, we identified core transcription factor differences between localized and metastatic tumors, thus providing insights into transcription factor networks that are critical for defining metastatic potential.

While our analysis revealed differential chromatin profiles between localized and primary metastatic, we were also able to identify altered chromatin activity in distal met tumors that showed enhanced Wnt signaling activity and signatures associated with cell migration, which as previously mentioned are consistent with metastatic osteosarcoma biology. Interestingly, we noted an enrichment in actin cytoskeletal reorganization in the primary met, but not in the local control or distal met tumors, which could indicate essential intrinsic structural alterations for these subsets of tumors. Future studies will focus on the identification of the epigenetic landscape associated with therapeutic resistant disease through analysis of additional patient-derived specimens as well as additional functional genomic studies on candidate metastatic-promoting genes, including PREX1, PPP1BD and IGF2B1P1. Finally, these genes can potentially be biomarkers of disease progression, or relapse, and eventually utilized to stratify patients into high versus low-risk disease.

## Acknowledgments

IS was supported by 1R21NS121945 and CPRIT RP230204. JTY was supported by 1R01EB026453, 1R01 CA21554 and 1R21CA267914 and The Faris D. Virani Ewing Sarcoma Center. PDX generation was supported by the CPRIT Core Facility Award (RP170691). We also acknowledge GyoungEun Kim for assistance in the generation of the PDXs used in this study.

## REFERNCES

1. Mirabello L, Troisi RJ, Savage SA. Osteosarcoma incidence and survival rates from 1973 to 2004: data from the Surveillance, Epidemiology, and End Results Program. Cancer: Interdisciplinary International Journal of the American Cancer Society 2009;115:1531–43

2. Duchman KR, Gao Y, Miller BJ. Prognostic factors for survival in patients with high-grade osteosarcoma using the Surveillance, Epidemiology, and End Results (SEER) Program database. Cancer epidemiology 2015;39:593–9

3. Jaffe N, Farber S, Traggis D, Geiser C, Kim BS, Das L, et al. Favorable response of metastatic osteogenic sarcoma to pulse high-dose methotrexate with citrovorum rescue and radiation therapy. Cancer 1973;31:1367–73

4. Heintzman ND, Hon GC, Hawkins RD, Kheradpour P, Stark A, Harp LF, et al. Histone modifications at human enhancers reflect global cell-type-specific gene expression. Nature 2009;459:108–12

5. Gifford CA, Ziller MJ, Gu H, Trapnell C, Donaghey J, Tsankov A, et al. Transcriptional and epigenetic dynamics during specification of human embryonic stem cells. Cell 2013;153:1149–63

6. Creyghton MP, Cheng AW, Welstead GG, Kooistra T, Carey BW, Steine EJ, et al. Histone H3K27ac separates active from poised enhancers and predicts developmental state. Proceedings of the National Academy of Sciences 2010;107:21931–6

7. Panigrahi A, O’Malley BW. Mechanisms of enhancer action: the known and the unknown. Genome biology 2021;22:1–30

8. Mack* SC, Singh* I, Wang* X, Hirsch R, Wu Q, Villagomez R, et al. Chromatin landscapes reveal developmentally encoded transcriptional states that define human glioblastoma. J Exp Med 2019;216:1071–90

9. Akhtar-Zaidi B, Cowper-Sal·lari R, Corradin O, Saiakhova A, Bartels CF, Balasubramanian D, et al. Epigenomic enhancer profiling defines a signature of colon cancer. Science 2012;336:736–9

10. Della Chiara G, Gervasoni F, Fakiola M, Godano C, D’Oria C, Azzolin L, et al. Epigenomic landscape of human colorectal cancer unveils an aberrant core of pan-cancer enhancers orchestrated by YAP/TAZ. Nature communications 2021;12:1–18

11. McDonald OG, Li X, Saunders T, Tryggvadottir R, Mentch SJ, Warmoes MO, et al. Epigenomic reprogramming during pancreatic cancer progression links anabolic glucose metabolism to distant metastasis. Nature genetics 2017;49:367–76

12. Kron KJ, Bailey SD, Lupien M. Enhancer alterations in cancer: a source for a cell identity crisis. Genome medicine 2014;6:1–12

13. Roe J-S, Hwang C-I, Somerville TD, Milazzo JP, Lee EJ, Da Silva B, et al. Enhancer reprogramming promotes pancreatic cancer metastasis. Cell 2017;170:875–88. e20

14. Morrow JJ, Bayles I, Funnell AP, Miller TE, Saiakhova A, Lizardo MM, et al. Positively selected enhancer elements endow osteosarcoma cells with metastatic competence. Nature medicine 2018;24:176–85

15. Rainusso N, Cleveland H, Hernandez JA, Quintanilla NM, Hicks J, Vasudevan S, et al. Generation of patient-derived tumor xenografts from percutaneous tumor biopsies in children with bone sarcomas. Pediatric Blood & Cancer 2019;66:e27579

16. Langmead B, Salzberg SL. Fast gapped-read alignment with Bowtie 2. Nature methods 2012;9:357–9

17. Zhang Y, Liu T, Meyer CA, Eeckhoute J, Johnson DS, Bernstein BE, et al. Model-based analysis of ChIP-Seq (MACS). Genome biology 2008;9:1–9

18. Kluin RJC, Kemper K, Kuilman T, de Ruiter JR, Iyer V, Forment JV, et al. XenofilteR: computational deconvolution of mouse and human reads in tumor xenograft sequence data. BMC Bioinformatics 2018;19:366

19. Akalin A, Franke V, Vlahoviček K, Mason CE, Schübeler D. Genomation: a toolkit to summarize, annotate and visualize genomic intervals. Bioinformatics 2015;31:1127–9

20. Dobin A, Davis CA, Schlesinger F, Drenkow J, Zaleski C, Jha S, et al. STAR: ultrafast universal RNA-seq aligner. Bioinformatics 2013;29:15–21

21. Tsherniak A, Vazquez F, Montgomery PG, Weir BA, Kryukov G, Cowley GS, et al. Defining a cancer dependency map. Cell 2017;170:564–76. e16

22. Wu T, Hu E, Xu S, Chen M, Guo P, Dai Z, et al. clusterProfiler 4.0: A universal enrichment tool for interpreting omics data. The Innovation 2021;2:100141

23. Maher B. ENCODE: The human encyclopaedia. Nature 2012;489:46–8

24. Anders S, Huber W. Differential expression analysis for sequence count data. Genome Biol 2010;11:R106

25. Belkhiri A, Zhu S, Chen Z, Soutto M, El-Rifai W. Resistance to TRAIL Is Mediated by DARPP-32 in Gastric CancerDARPP-32 Mediates Resistance to TRAIL. Clinical cancer research 2012;18:3889–900

26. Christenson JL, Denny EC, Kane SE. t-Darpp overexpression in HER2-positive breast cancer confers a survival advantage in lapatinib. Oncotarget 2015;6:33134

27. Gu L, Waliany S, Kane SE. Darpp-32 and its truncated variant t-Darpp have antagonistic effects on breast cancer cell growth and herceptin resistance. PloS one 2009;4:e6220

28. Vangamudi B, Peng D-F, Cai Q, El-Rifai W, Zheng W, Belkhiri A. t-DARPP regulates phosphatidylinositol-3-kinase-dependent cell growth in breast cancer. Molecular cancer 2010;9:1–11

29. Wang M, Pan Y, Liu N, Guo C, Hong L, Fan D. Overexpression of DARPP-32 in colorectal Adenocarcinoma. International journal of clinical practice 2005;59:58–61

30. Beckler A, Moskaluk CA, Zaika A, Hampton GM, Powell SM, Frierson Jr HF, et al. Overexpression of the 32-kilodalton dopamine and cyclic adenosine 3L, 5L-monophosphate-regulated phosphoprotein in common adenocarcinomas. Cancer: Interdisciplinary International Journal of the American Cancer Society 2003;98:1547–51

31. Belkhiri A, Zhu S, El-Rifai W. DARPP-32: from neurotransmission to cancer. Oncotarget 2016;7:17631

32. Alam S, Astone M, Liu P, Hall SR, Coyle AM, Dankert EN, et al. DARPP-32 and t-DARPP promote non-small cell lung cancer growth through regulation of IKKα-dependent cell migration. Communications biology 2018;1:1–15

33. Huang X, Zhang H, Guo X, Zhu Z, Cai H, Kong X. Insulin-like growth factor 2 mRNA-binding protein 1 (IGF2BP1) in cancer. Journal of hematology & oncology 2018;11:1–15

34. Wang L, Aireti A, Aihaiti A, Li K. Expression of microRNA-150 and its target gene IGF2BP1 in human osteosarcoma and their clinical implications. Pathology & Oncology Research 2019;25:527–33

35. Marei H, Carpy A, Woroniuk A, Vennin C, White G, Timpson P, et al. Differential Rac1 signalling by guanine nucleotide exchange factors implicates FLII in regulating Rac1-driven cell migration. Nature communications 2016;7:1–16

36. Baker MJ, Abba MC, Garcia-Mata R, Kazanietz MG. P-REX1-independent, calcium-dependent RAC1 hyperactivation in prostate cancer. Cancers 2020;12:480

37. Welch HC, Coadwell WJ, Ellson CD, Ferguson GJ, Andrews SR, Erdjument-Bromage H, et al. P-Rex1, a PtdIns (3, 4, 5) P3-and Gβγ-regulated guanine-nucleotide exchange factor for Rac. Cell 2002;108:809–21

38. Lindsay CR, Lawn S, Campbell AD, Faller WJ, Rambow F, Mort RL, et al. P-Rex1 is required for efficient melanoblast migration and melanoma metastasis. Nature communications 2011;2:1–9

39. Sosa MS, Lopez-Haber C, Yang C, Wang H, Lemmon MA, Busillo JM, et al. Identification of the Rac-GEF P-Rex1 as an essential mediator of ErbB signaling in breast cancer. Molecular cell 2010;40:877–92

40. Srijakotre N, Liu H-J, Nobis M, Man J, Yip HYK, Papa A, et al. PtdIns (3, 4, 5) P3-dependent Rac exchanger 1 (P-Rex1) promotes mammary tumor initiation and metastasis. Proceedings of the National Academy of Sciences 2020;117:28056–67

41. Heinz S, Benner C, Spann N, Bertolino E, Lin YC, Laslo P, et al. Simple combinations of lineage-determining transcription factors prime cis-regulatory elements required for macrophage and B cell identities. Molecular cell 2010;38:576–89

42. Li M, Shen Y, Wang Q, Zhou X. MiR-204-5p promotes apoptosis and inhibits migration of osteosarcoma via targeting EBF2. Biochimie 2019;158:224–32

43. Patiño-García A, Zalacain M, Folio C, Zandueta C, Sierrasesúmaga L, San Julián M, et al. Profiling of Chemonaive Osteosarcoma and Paired-Normal Cells Identifies EBF2 as a Mediator of Osteoprotegerin Inhibition to Tumor Necrosis Factor–Related Apoptosis-Inducing Ligand–Induced Apoptosis. Clinical Cancer Research 2009;15:5082–91

44. Goel S, Bhatia V, Kundu S, Biswas T, Carskadon S, Gupta N, et al. Transcriptional network involving ERG and AR orchestrates Distal-less homeobox-1 mediated prostate cancer progression. Nature communications 2021;12:1–22

45. Rahman NI, Abdul Murad NA, Mollah MM, Jamal R, Harun R. NFIX as a master regulator for lung cancer progression. Frontiers in pharmacology 2017;8:540

46. Zhang Y, Wang X. Targeting the Wnt/β-catenin signaling pathway in cancer. Journal of hematology & oncology 2020;13:1–16

47. Migliardi G, Sassi F, Torti D, Galimi F, Zanella ER, Buscarino M, et al. Inhibition of MEK and PI3K/mTOR suppresses tumor growth but does not cause tumor regression in patient-derived xenografts of RAS-mutant colorectal carcinomas. Clin Cancer Res 2012;18:2515–25

48. Gao H, Korn JM, Ferretti S, Monahan JE, Wang Y, Singh M, et al. High-throughput screening using patient-derived tumor xenografts to predict clinical trial drug response. Nat Med 2015;21:1318–25

49. Bertotti A, Migliardi G, Galimi F, Sassi F, Torti D, Isella C, et al. A molecularly annotated platform of patient-derived xenografts (“xenopatients”) identifies HER2 as an effective therapeutic target in cetuximab-resistant colorectal cancer. Cancer Discov 2011;1:508–23

50. Lazzari L, Corti G, Picco G, Isella C, Montone M, Arcella P, et al. Patient-Derived Xenografts and Matched Cell Lines Identify Pharmacogenomic Vulnerabilities in Colorectal Cancer. Clin Cancer Res 2019;25:6243–59

51. Bleijs M, van de Wetering M, Clevers H, Drost J. Xenograft and organoid model systems in cancer research. EMBO J 2019;38:e101654

52. Khan A, Mathelier A. Intervene: a tool for intersection and visualization of multiple gene or genomic region sets. BMC bioinformatics 2017;18:1–8

