## Supplemental Figures for "Intrinsic epigenetic state of primary osteosarcoma drives metastasis"

**Running title:** Intrinsic epigenetic state of primary OS drives metastasis

**Keywords:** Osteosarcoma; Epigenetics; ChIP-Seq; ATAC-seq; Metastasis.

**Financial support:** IS was supported by 1R21NS121945 and CPRIT RP230204**.** JTY was supported by 1R01EB026453, 1R01 CA21554 and 1R21CA267914 and The Faris D. Virani Ewing Sarcoma Center. PDX Core was supported by the CPRIT Core Facilities Support Grant RP170691.

**Co-corresponding authors**: Dr. Irtisha Singh, Department of Cell Biology and Genetics, College of Medicine, Texas A&M University, 8447 Riverside Pkwy MREB II, Suite 4344, Bryan, TX 77807-3260; Dr. Jason Yustein, Emory University, Health Sciences Research Building, 1760 Haygood Drive, Atlanta, GA 30322

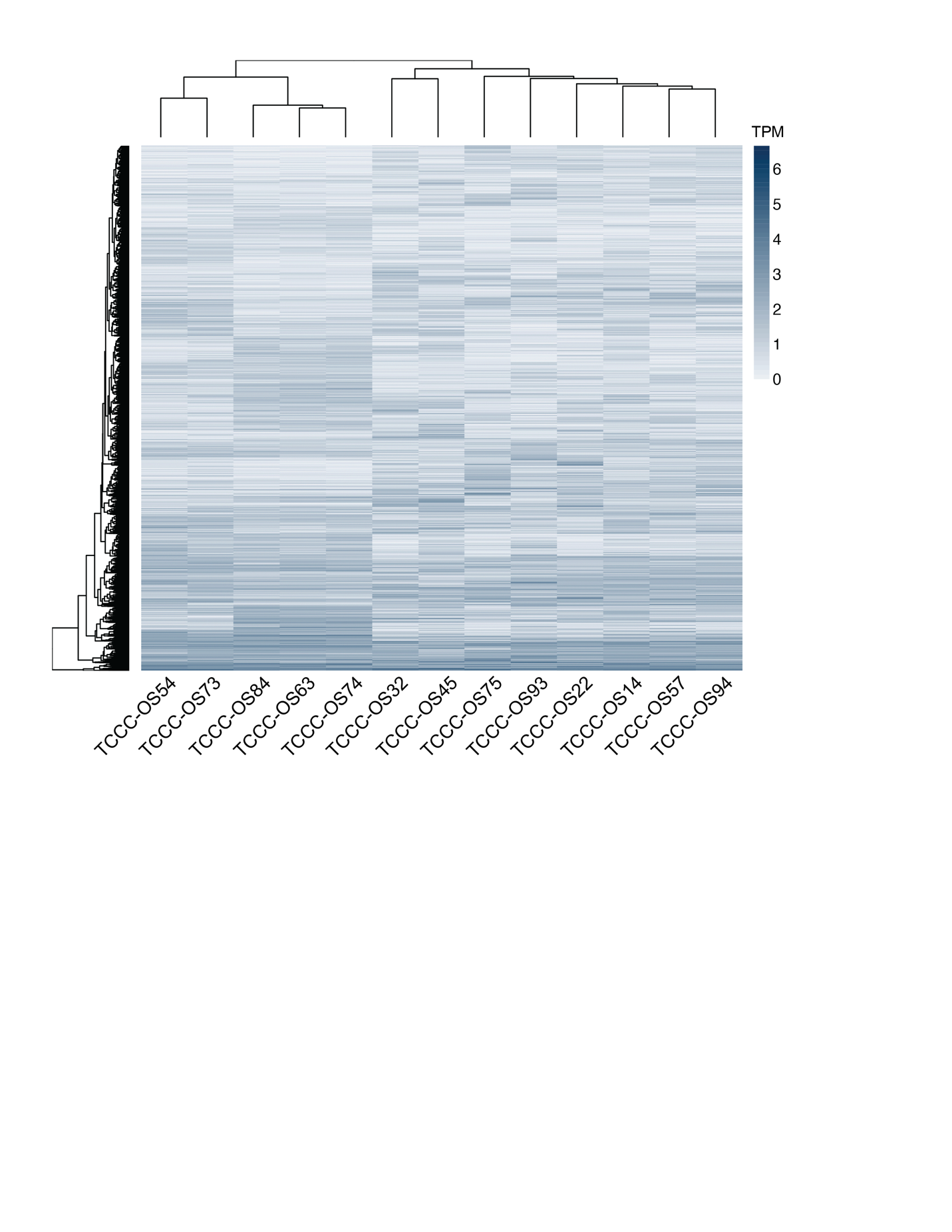

**Supplementary S1 – Hierarchical clustering of signal intensity (tags per million) of top 20% most variable regions by median absolute deviation (n = 31,690) of all H3K27ac enriched peaks.**

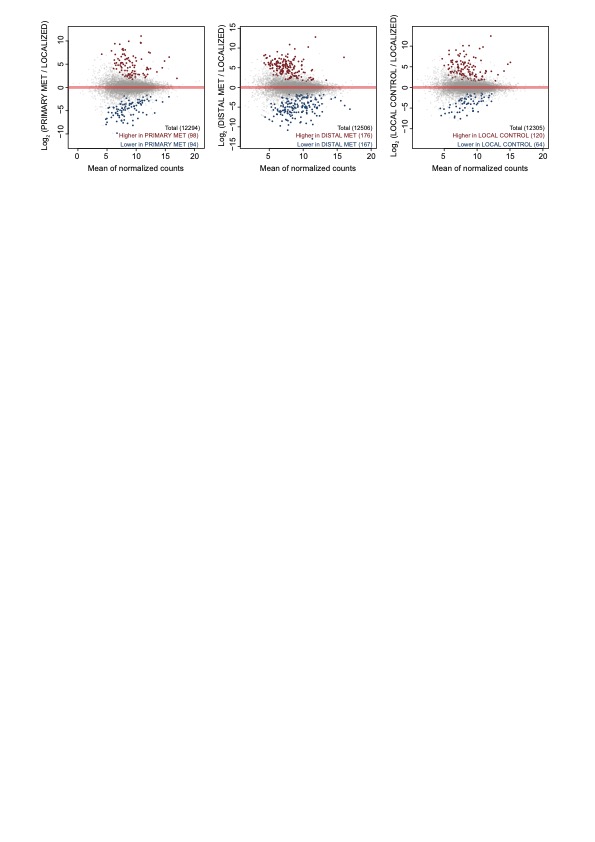

**Supplementary S2 – Differential expression of RNA-seq between the different groups**.

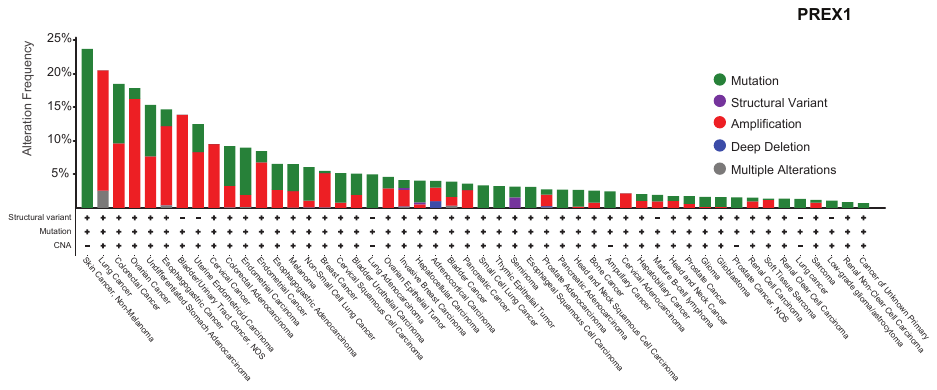

**Supplementary S3– Gene alteration frequency of PREX1**. Gene alteration frequency obtained from cBio portal.

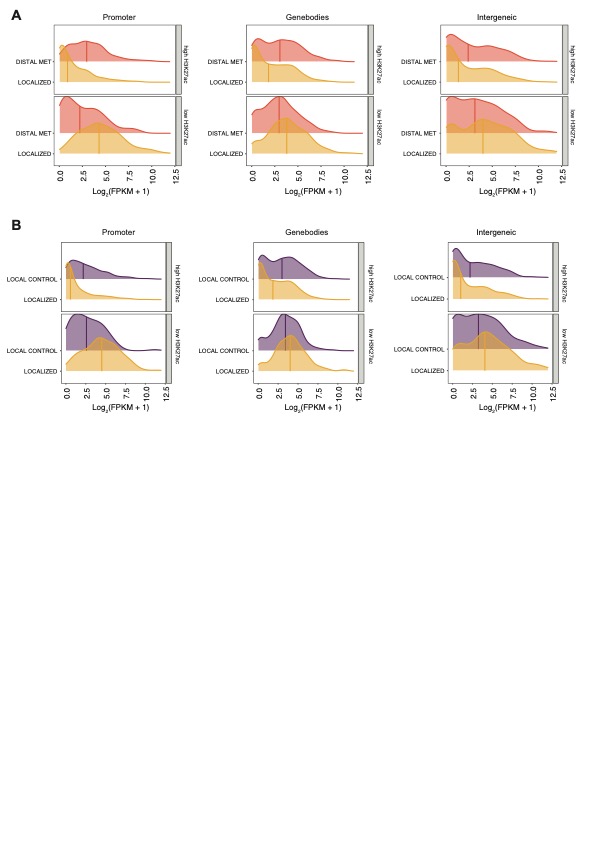

**Supplementary S4** – **Differential chromatin activity shapes transcriptional profile** **(A)** Same as Fig 2E but for the comparison between distal met and localized tumors **(B)** Same as Fig 2E but for the comparison between local control and localized tumors.

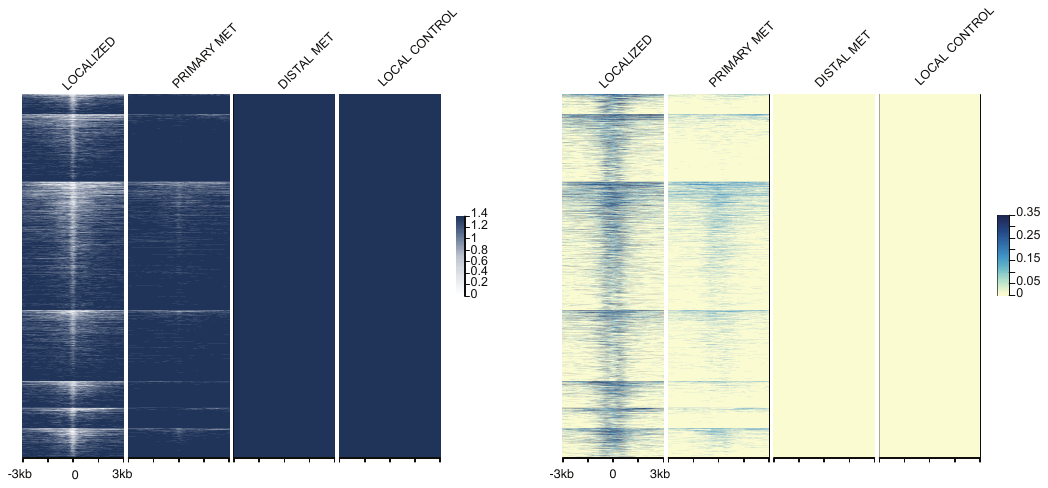

**Supplementary S5 – Genomic regions with reduced chromatin activity.** Same as 7A but for regions with reduced chromatin activity.
